# Redistribution of a glucuronoxylomannan epitope towards the capsule surface coincides with Titanisation in the human fungal pathogen *Cryptococcus neoformans*

**DOI:** 10.1101/431650

**Authors:** Mark Probert, Xin Zhou, Margaret Goodall, Simon A. Johnston, Ewa Bielska, Elizabeth R. Ballou, Robin C. May

## Abstract

Disseminated infections with the fungal species *Cryptococcus neoformans* or, less frequently, *C. gattii,* are a leading cause of mortality in immunocompromised individuals. Central to the virulence of both species is an elaborate polysaccharide capsule that consists predominantly of glucuronoxylomannan (GXM). Due to its abundance, GXM is an ideal target for host antibodies, and several monoclonal antibodies (mAbs) have previously been derived using purified GXM or whole capsular preparations as antigen. In addition to their application in the diagnosis of cryptococcosis, anti-GXM mAbs are invaluable tools for studying capsule structure. In this study, we report the production and characterisation of a novel anti-GXM mAb, Crp127, that unexpectedly reveals a role for GXM remodelling during the process of fungal Titanisation. We show that Crp127 recognises a GXM epitope in an *O*-acetylation dependent, but xylosylation-independent, manner. The epitope is differentially expressed by the four main serotypes of *Cryptococcus neoformans* and *gattii,* is heterogeneously expressed within clonal populations of *C. gattii* serotype B strains and is typically confined to the central region of the enlarged capsule. Uniquely, however, this epitope redistributes to the capsular surface in Titan cells, a recently recognised subset of giant fungal cells that are produced in the host lung and are critical for successful infection. Crp127 therefore highlights hitherto unexpected features of cryptococcal morphological change and may hold significant therapeutic potential in differentially identifying cryptococcal strains and subtypes.

**Importance:** *Cryptococcus neoformans* and *Cryptococcus gattii* are the etiological agents of cryptococcosis, an invasive fungal infection responsible for approximately 200,000 deaths each year and 15% of AIDS-related deaths annually. Whilst the main virulence factor for both species is a highly variable polysaccharide capsule, formation of Titan cells also underlies the pathogenesis of *C. neoformans.* Previous studies have shown that capsule composition differs between yeast and Titan cells, however no clear distinctions in the expression or localisation of specific capsular epitopes have been made. In this study, we characterise a novel monoclonal antibody (mAb) specific to a capsular epitope that is differentially distributed throughout the capsules produced by yeast and Titan cells. Whilst this epitope is found within the midzone of yeast capsules, the presentation of this epitope on the surface of Titan cell capsules may represent a way in which these cell types are perceived differently by the immune system.

## Introduction

As the two main etiological agents of cryptococcosis, *Cryptococcus neoformans* and *Cryptococcus gattii* are major contributors to the global health burden imposed by invasive fungal infections (1). Whilst *C. neoformans* typically manifests as meningitis in immunocompromised individuals, *C. gattii* infections are not associated with specific immune defects and have been responsible for fatal outbreaks of pneumonia (2–4). Central to the virulence of both species is an elaborate polysaccharide capsule, without which *Cryptococcus* is rendered avirulent (5, 6). The composition of this capsule is highly variable and differs between yeast cells and Titan cells, the latter representing a subset of giant cells (>10 μm in cell body diameter) formed by *C. neoformans* within the host lung (7–9). Titan cells contribute to pathogenesis by resisting phagocytosis, enhancing dissemination of yeast to the central nervous system and altering host immune status (7, 9–13).

The cryptococcal capsule consists of ~90% glucuronoxylomannan (GXM), ~10% glucuronoxylomannogalactan (GXMGal) and <1% mannoproteins (MPs) (14). GXM is a megadalton polysaccharide containing a backbone of a-(1,3)-mannan that is decorated with β-(1,2)-glucuronic acid, β-(1,2)-xylose and β-(1,4)-xylose substituents (15). The backbone mannan can also be *O*-acetylated, although the position at which this modification is added remains unclear for most strains (14–16). Seven repeat motifs - called structure reporter groups (SRGs) - contribute to structural variation in GXM (15). All SRGs contain a β-(1,2)-glucuronic acid on their first mannose residue, however the number of β-(1,2)- and β-(1,4)-xylose substituents varies (15). The extent and position of *O*-acetyl groups in each SRG remain unclear, however xylose and *O*-acetyl groups attached to the same mannose residue appear to be mutually exclusive (17). SRG usage differs between the four main serotypes of *Cryptococcus,* with each strain designated a serotype based on the reactivity of its capsular material with antibody preparations (18). *C. neoformans* serotypes A and D tend to biosynthesise GXM containing SRGs with fewer xylose substituents than those from *C. gattii* serotypes B and C (15, 19).

Whilst capsule structure differs between serotypes of *Cryptococcus,* a flexible biosynthetic pathway enables rapid remodelling of the capsule under different environmental conditions (20). *In vitro*, changes in *O*-acetylation have been associated with cell ageing in *C. neoformans* (21), reaffirming earlier reports that capsules produced within clonal populations are far from homogeneous (19, 22). *In vivo*, changes in capsule size and structure coincide with the infection of different organs and likely enhance fitness through the evasion of host immunity (23–25). In light of these observations, it is perhaps unsurprising that capsules produced by Titan cells are structurally distinct from those produced by typical yeast cells (7, 11, 26). As the increased chitin content of cell walls produced by Titan cells is associated with activation of a detrimental TH2 immune response during cryptococcosis (27), it is possible that hitherto unidentified structural differences in Titan cell capsules also contribute to the modulation of host immunity by this *C. neoformans* morphotype.

Alterations in capsule structure are likely to affect how *Cryptococcus* is perceived by host immune molecules, with antibodies particularly sensitive to small changes in molecular structures. Following exposure to cryptococci, immunoglobulin M (IgM) antibodies are the most abundant isotype of antibody produced in response to GXM (28). As a repetitive capsular polysaccharide, GXM is a T-independent type 2 antigen and antibodies generated against it utilise a restricted set of variable region gene segments (29). Using monoclonal antibodies (mAbs) in conjunction with mutants harbouring specific defects in GXM modification (17, 30, 31), it has been determined that *O*-acetylation and, to a lesser extent, xylosylation of GXM are important for epitope recognition by anti-GXM antibodies (16, 30). Whilst there is no consensus surrounding the effect of GXM *O*-acetylation on virulence (17, 32), its influence on antibody binding suggests that changes in GXM *O*-acetylation could be a strategy deployed by cryptococci to avoid recognition by immune effectors. Additionally, despite the immunomodulatory roles for GXM *O*-acetylation that have been identified (30, 33), receptors that bind *O*-acetylated GXM remain elusive (34). Due to the enigmatic nature of this modification within the primary virulence factor of cryptococci, further investigation of GXM *O*-acetylation will help unravel the complexities of cryptococcal capsule structure with the ultimate aim of understanding the strategies deployed by this fatal fungal pathogen to evade host immunity.

In the present study, we report the generation of Crp127, a murine IgM mAb, using a cocktail of heat-killed *C. neoformans* H99 (serotype A), heat-killed *C. gattii* R265 (serotype B) and their lysates as an immunogen. Characterisation of Crp127 demonstrated that it is an *O*-acetyl-dependent anti-GXM mAb specific to an epitope expressed by the four *Cryptococcus* serotypes in a serotype-specific manner. Having subsequently found that this epitope is heterogeneously expressed within serotype B populations and is spatially confined to distinct regions of the enlarged capsule across all strains tested, we then turned our attention to its expression by Titan cells. Intriguingly, we noticed that the spatial distribution of this epitope differs within the capsules produced by the three *C. neoformans* morphotypes found within Titanising populations. Further analysis revealed that, under conditions permissive for Titanisation, cell enlargement coincides with the gradual redistribution of this epitope to the capsule surface.

## Results

### Crp127 recognises a capsular epitope located in GXM

During hybridoma screening, Crp127 was identified as staining the outer zone of live cryptococci. We first assessed whether Crp127 recognises a capsular component by performing flow cytometric analysis of three GXM-deficient mutants (R265 *cap10*Δ, Kn99α *cap59*Δ and B3501 *cap67*Δ), a GXMGal-deficient mutant (Kn99α *uge1*Δ) and a mutant lacking both GXM and GXMGal (Kn99α *cap59*Δ*uge1*Δ), using an Alexa-488-conjugated anti-IgM secondary antibody to label Crp127. Unlike their corresponding wild-type strains, the GXM-deficient mutants were not recognised by Crp127 (fig. 1A-C; *cap10*Δ P < 0.05; *cap67*Δ P < 0.01*; cap59*Δ P < 0.01; *cap59*Δ*uge1*Δ P < 0.01, Student’s t-test). In contrast, the GXMGal-deficient *uge1*Δ mutant was bound at levels similar to the wild-type strain (fig. 1C; P > 0.05). Confocal microscopy corroborated these observations, with no observable binding of Crp127 to GXM-deficient mutants but clear binding of Crp127 to the GXMGal-deficient mutant (fig. 1D-F). Taken together, these experiments demonstrated that the epitope recognised by Crp127 – hereon referred to as the Crp127 epitope – is a component of GXM.

**Fig. 1.**
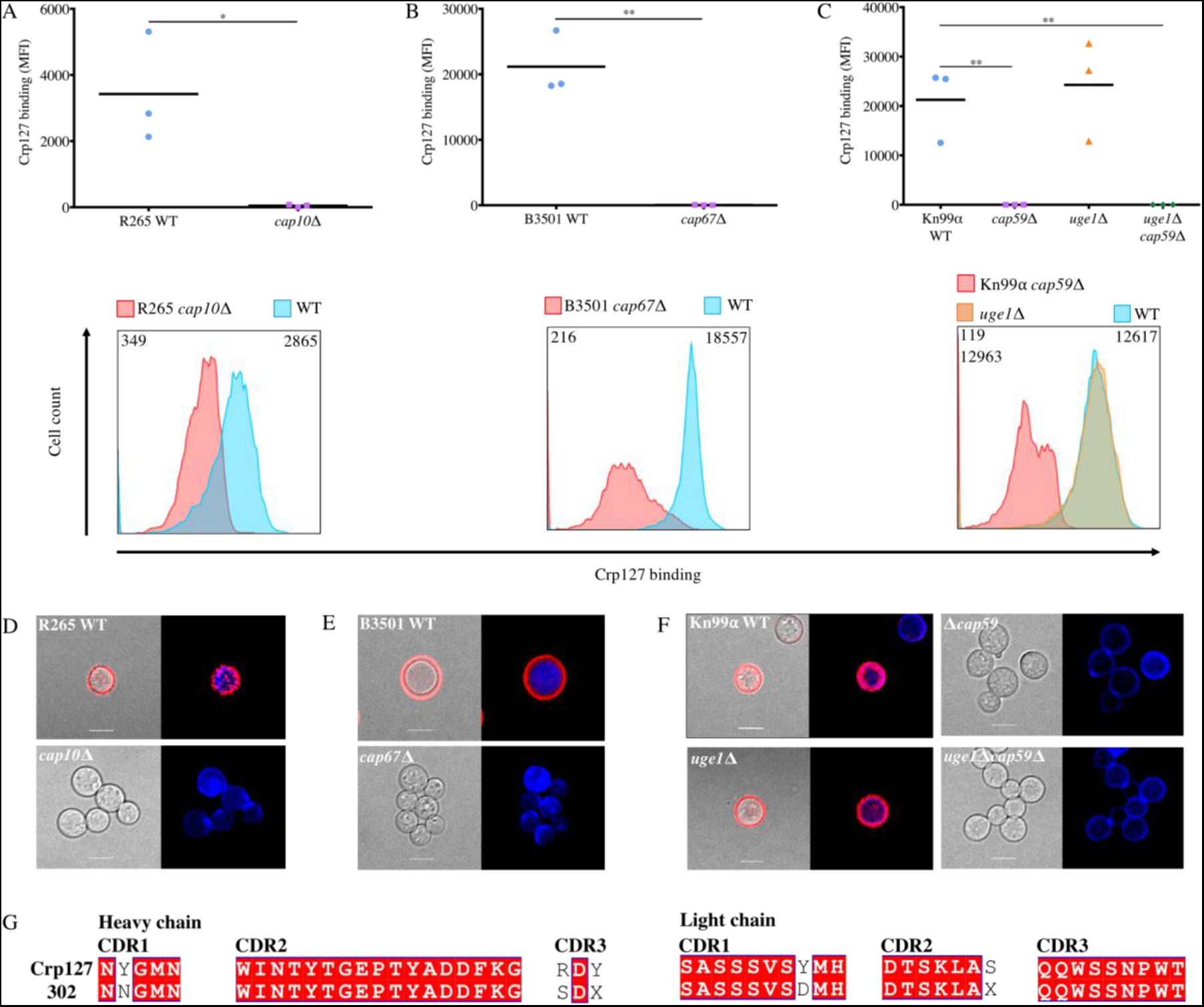
Crp127 is an anti-GXM mAb. The ability of Crp127 to bind to GXM- and GXMGal-deficient mutants of C. gattii and C. neoformans was quantified via flow cytometry. Scatter plots (top row) and representative histograms (bottom row) are presented for **A)** R265 cap10Δ, **B)** B3501 cap67Δ, **C)** Kn99α cap59Δ, Kn99α uge1Δ, Kn99α cap59Δuge1Δ and their corresponding wild-type strains. For scatter plots, corrected median fluorescence intensity (MFI) values were calculated by subtracting the MFI value of isotype control cells from the MFI value of the Crp127-treated cells, with data points representing MFI values calculated from three biological replicates performed as independent experiments. A Student’s t-test was used to test for statistically significant differences between R265 cap10Δ and B3501 cap67Δ and their corresponding wild-type strains, whilst one-way ANOVA followed by Dunnett’s multiple comparison test was used to test for statistically significant differences between Kn99α cap59Δ, Kn99α uge1Δcap59Δ, Kn99α uge1Δ and the wild-type strain KN99α (n=3) (ns P > 0.05; * P < 0.05; ** P < 0.01). Histograms show a representative distribution of Crp127 binding for one or all of the strains in the above scatter plot, with the colour-coded key provided for reference. Numerical values in the top left and right of each histogram correspond to the MFI value calculated from the strain labelled directly above. **D)** R265 cap10Δ, **E)** B3501 cap67Δ, **F)** Kn99α cap59Δ, uge1Δ, uge1Δcap59Δ and their wild-type strains were labelled for chitin using calcofluor-white (CFW; blue) and Crp127 (far-red; goat Alexa-647-conjugated anti-mouse IgM μ-chain) and maximum-intensity projections generated from confocal microscopy z-stacks. Presented are representative images merged for transmitted light and Crp127 (left panels) and Crp127 and chitin (right panels). Scale bars represent 5 μm. **G)** Amino acid sequences of the CDRs from mAbs Crp127 and 302 were aligned. Residues highlighted in red are identical. X denotes amino acid residues that are undetermined.

In order to draw comparisons with previously described anti-GXM mAbs, we next sequenced both the heavy-chain variable (V_H_) and light-chain variable (V_L_) regions of Crp127 (fig. S1). Antibody sequence analysis revealed that the Crp127 V_H_ region uses the V_H_9(VGAM family)-J_H_2 combination with a D-segment consisting of four amino acids, whilst the V_L_ region uses a combination of V_k_4/5-J_k_1. This gene usage profile is remarkably similar to that of anti-GXM IgG1 mAb 302, which differs only in respect to the length of its D-segment (35). In line with this, alignment of Crp127 and 302 protein sequences revealed 86.6% and 98.1% sequence identity between V_H_ and V_L_ regions, respectively (fig. S1A-B). When the complementarity determining regions (CDRs) of mAbs Crp127 and 302 were aligned, we found identical V_H_ CDR2 and V_L_ CDR3 regions whilst the remaining CDRs differed at one or two positions (fig. 1G). Notably, however, the sequence available for mAb 302 is incomplete and may contribute additional variation (fig. 1G). We also compared the CDRs of Crp127 with four other well-characterised anti-GXM IgM mAbs – 2D10, 12A1, 13F1 and 21D2 – with less similarities identified between Crp127 and these mAbs (fig. S1C-D).

### GXM *O*-acetylation is required for Crp127 epitope recognition

Considering the importance of *O*-acetylation and xylosylation to the antigenic signature of GXM (30), we proceeded to investigate the effect of these modifications on Crp127 epitope recognition. We firstly tested the ability of Crp127 to recognise two xylose-deficient mutants (JEC155 *uxs1*Δ (serotype D) and Kn99α *uxs1*Δ (serotype A)). No significant differences were found between either *uxs1*Δ mutant and their corresponding wild-type strains (fig. 2A; JEC155 *uxs1*Δ P > 0.05; Kn99α *uxs1*Δ P > 0.05), indicating that xylosylation does not impact Crp127 binding. In contrast, however, antibody binding was completely abrogated in the *O*-acetyl-deficient *cas1*Δ mutant (fig. 2B; P < 0.01), indicating that *O*-acetylation of GXM is an essential prerequisite for Crp127 epitope recognition.

Having made this observation, we proceeded to test two further mutants in genes implicated in GXM *O*-acetylation. Kn99α *cas3*Δ exhibits a ~70% reduction in GXM *O*-acetylation, whereas Kn99α *cas31*Δ exhibits subtle differences in sugar composition of GXM but no reduction in GXM *O*-acetylation (36). Binding of Crp127 to the *cas3*Δ mutant was slightly reduced but statistically to the wild-type strain (fig. 2C; P > 0.05). This may reflect reduced density of *O*-acetylation in GXM produced by this mutant Surprisingly, however, Crp127 completely failed to recognise the *cas31*Δ mutant despite this strain retaining an *O*-acetylation profile similar to the wild-type (36) (fig. 2C; P < 0.01). To be certain that the *O*-acetyl-defective mutants tested still produced capsule, we confirmed the binding of *O*-acetyl-independent anti-GXM mAb F12D2 to each strain (fig. 2E-F; insets). Thus, *CAS1* and *CAS31* contribute to the formation of an *O*-acetylation dependent Crp127 epitope.

**Fig. 2.**
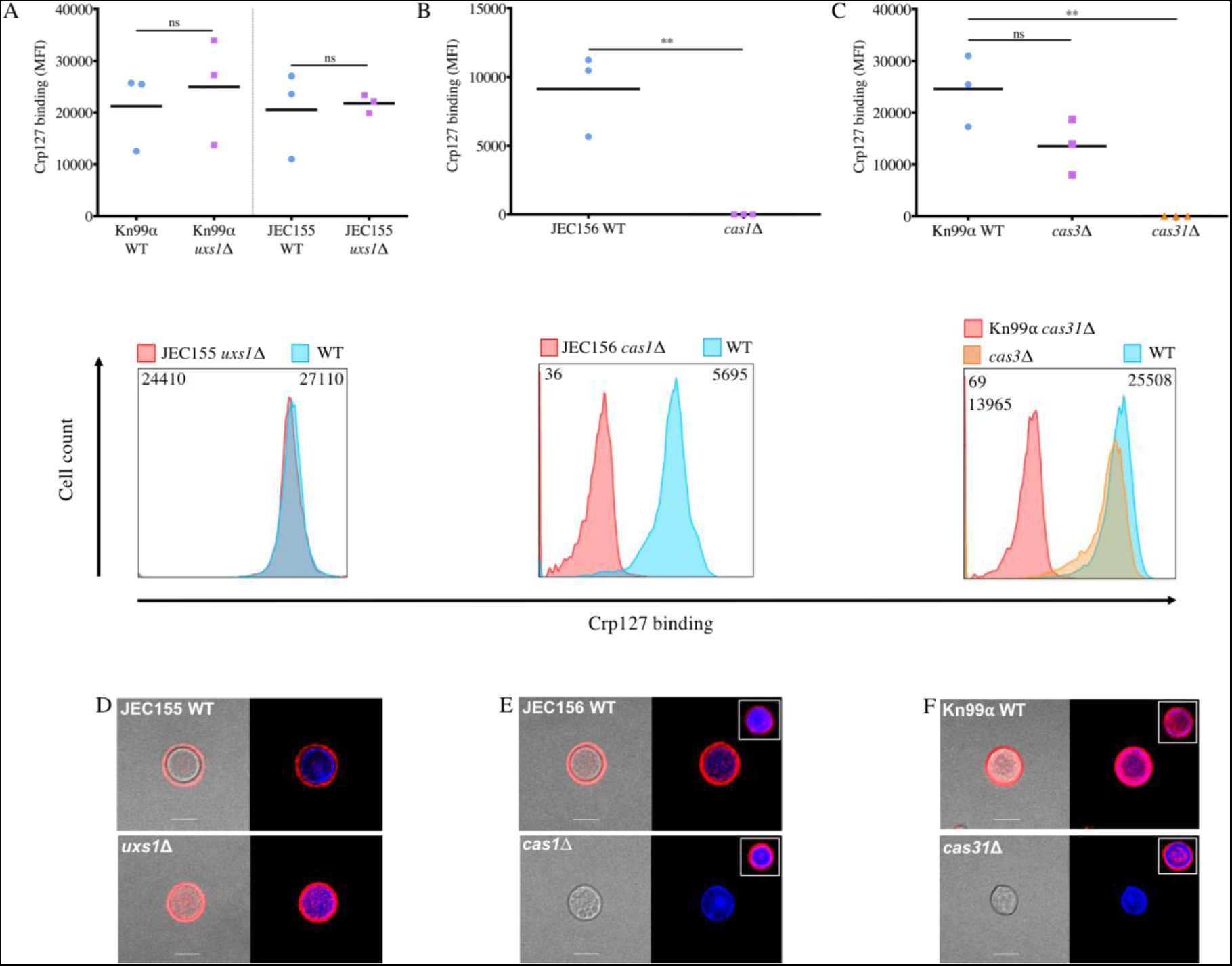
Crp127 requires O-acetylation, but not xylosylation, of GXM for epitope recognition. The ability of Crp127 to recognise mutants with specific defects in GXM modification was quantified via flow cytometry. Scatter plots (top row) and representative histograms (bottom row) are presented for **A)** Kn99α uxs1Δ, JEC155 uxs1Δ and corresponding wild-type strains; **B)** JEC156 cas1Δ and JEC155 wild-type and **C)** Kn99α cas3Δ, Kn99α cas31Δ and Kn99α wild-type. For scatter plots, corrected MFI values were calculated by subtracting the MFI value of isotype control cells from the MFI value of the Crp127-treated cells, with data points representing MFI values calculated from three biological replicates performed as independent experiments. A Student’s t-test was used to test for statistically significant differences between Kn99α uxs1Δ, JEC155 uxs1Δ and JEC156 cas1Δ and their corresponding wild-type strain whilst Dunnet’s multiple comparison test was used to test for statistically significant differences between Kn99α Δcas3, Kn99α Δcas31 mutants and the Kn99α wild-type strain (n=3) (ns P > 0.05; * P < 0.05; ** P < 0.01). Histograms show a representative distribution of Crp127 binding for one or all of the strains in the above scatter plot, with a colour-coded key provided for reference. Numerical values in the top left and right of each histogram correspond to the MFI value calculated from the strain labelled directly above. **D)** JEC155 uxs1Δ, **E)** JEC156 cas1Δ, **F)** Kn99α cas31Δ and corresponding wild-type strains were labelled for chitin (blue; CFW) and Crp127 (far-red; goat Alexa-647-conjugated goat anti-mouse IgM μ-chain) and maximum-intensity projections generated via confocal microscopy. Presented are representative images merged for transmitted light and Crp127 (left panels) and Crp127 and chitin (right panels). Insets in the top right of images show representative cells labelled for chitin (blue) and O-acetyl-independent mAb F12D2 (far-red; Alexa-647-conjugated F(abߣ)2 goat anti-Mouse IgG (H+L)) used as a control. Scale bars represent 5 μm.

Lastly, we confirmed that GXM *O*-acetylation is essential for Crp127 recognition by testing the binding of Crp127 to *Cryptococcus* cells that had been chemically de-*O*-acetylated via alkaline hydrolysis. For both strains tested (H99 and B3501), Crp127 failed to bind chemically de-*O*-acetylated cells, just as it failed to recognise the *cas1*Δ mutant (fig. S2A-C; H99 P < 0.01 and B3501 P < 0.01). However, binding of *O*-acetylation independent antibodies F12D2 and 18B7 remained unaltered (fig. S2C-D; P > 0.05). Taken together, our results demonstrate an essential role for *O*-acetylation in recognition of GXM by Crp127.

### *Cryptococcus* serotypes differ in their level of Crp127 epitope recognition

Differences in the *O*-acetylation state of GXM contributes towards serotype classification and is a source of structural variation within the capsule of cells from a clonal population (21, 30). Therefore, with Crp127 recognising an *O*-acetyl-dependent epitope, we next checked for differences in Crp127 staining between the five recognised serotypes of *Cryptococcus neoformans* and *gattii*, testing two independent strains of each serotype. Flow cytometry analysis demonstrated that Crp127 consistently bound most effectively to serotype D strains (B3501 and JEC155) (fig. 3A-B), with all cells within these populations exhibiting high-level accessibility of the Crp127 epitope (fig. 3F). We detected slightly lower binding to serotype A strains (fig. 3A-B), with high-level homogeneous staining also seen in the case of H99, but a proportion of unstained cells from strain Kn99α (fig. 3C). Interesting, the two AD hybrid strains tested (CBS 950 and ZG287) were notably different in regard to Crp127 binding (fig. 3A-B), with CBS 950 exhibiting low-level heterogeneous staining and ZG287 showing high-level homogeneous staining (fig. 3G).

The two remaining cryptococcal serotypes, B and C, together represent *C. gattii*. Serotype B strains R265 and CDCR272 demonstrated significantly lower epitope recognition than *C. neoformans* serotypes (fig. 3A-B) and considerable heterogeneity within the population (fig. 3D). Interestingly, however, serotype C strains were completely unrecognised by Crp127, with neither strain CBS 10101 or M27055 showing detectable staining (fig. 3A, B and E). From this, we conclude that there are serotype-specific differences in the availability of the Crp127 epitope, with epitope accessibility being related to serotype in a pattern of D > A >> B >>> C.

**Fig. 3.**
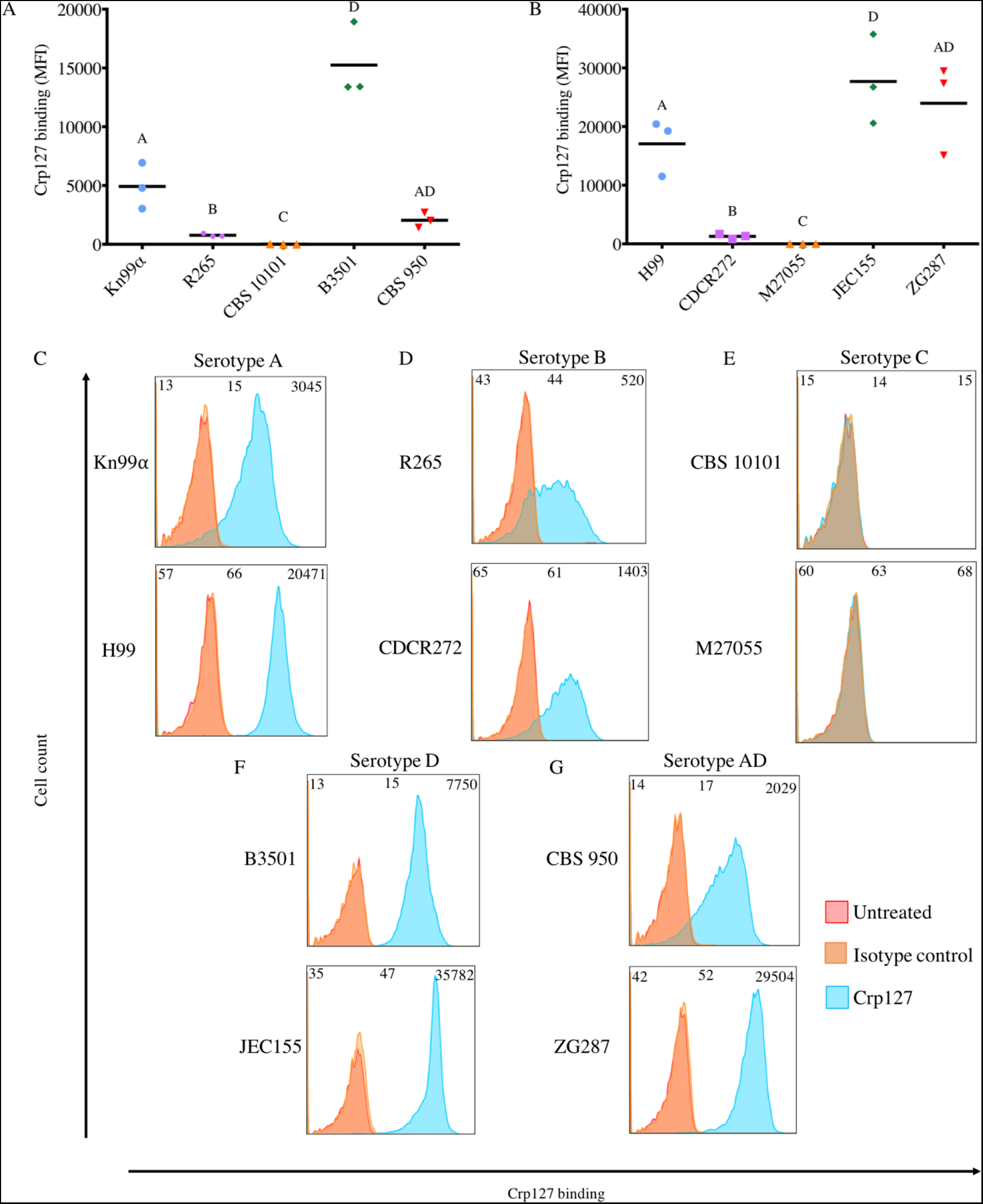
Recognition levels of the Crp127 epitope are associated with serotype. The ability of Crp127 to bind to two different strains from each Cryptococcus serotypes A, B, C, D and AD was quantified using flow cytometry. **A-B)** Scatter plots show corrected MFI values for each strain, which were calculated by subtracting the MFI value of isotype control cells from the MFI value of the corresponding Crp127-treated cells. Data points represent MFI values calculated from three biological replicates performed as independent experiments (n=3). Histograms show a representative distribution of Crp127 binding for **C)** serotype A strains Kn99α and H99, **D)** serotype B strains R265 and CDCR272, **E)** serotype C strains CBS 10101 and M27055, **F)** serotype D strains B3501 and JEC155 and **G)** serotype AD hybrid strains CBS 950 and ZG287. Fluorescence intensity values for untreated, isotype control and Crp127-treated cells are presented in red, blue and orange, respectively, with corresponding MFI values displayed in the top left, centre and right of each panel.

### Crp127 exhibits serotype-specific binding patterns that are not associated with opsonic efficacy

Having identified differential levels of the Crp127 epitope between serotypes using flow cytometry, we next examined their patterns of binding by immunofluorescence microscopy. Indirect immunofluorescence revealed an annular binding pattern for all four strains representing serotypes A and D (fig. 4A and D). In line with their differences in flow cytometry, the two AD hybrid strains tested showed different patterns of binding, with CBS 950 showing punctate binding and ZG287 showing a mix of annular and punctate staining. Both *C. gattii* serotype B strains exhibited punctate binding (fig. 4B and E) whilst, in agreement with flow cytometry, no Crp127 binding was detected when imaging serotype C strains CBS 10101 or M27055. However, *O-*acetyl-independent mAb F12D2 bound well to these strains, suggesting that the lack of Crp127 binding reflects changes in GXM *O*-acetylation rather than loss of capsular material (fig. 4C; inset).

**Fig. 4.**
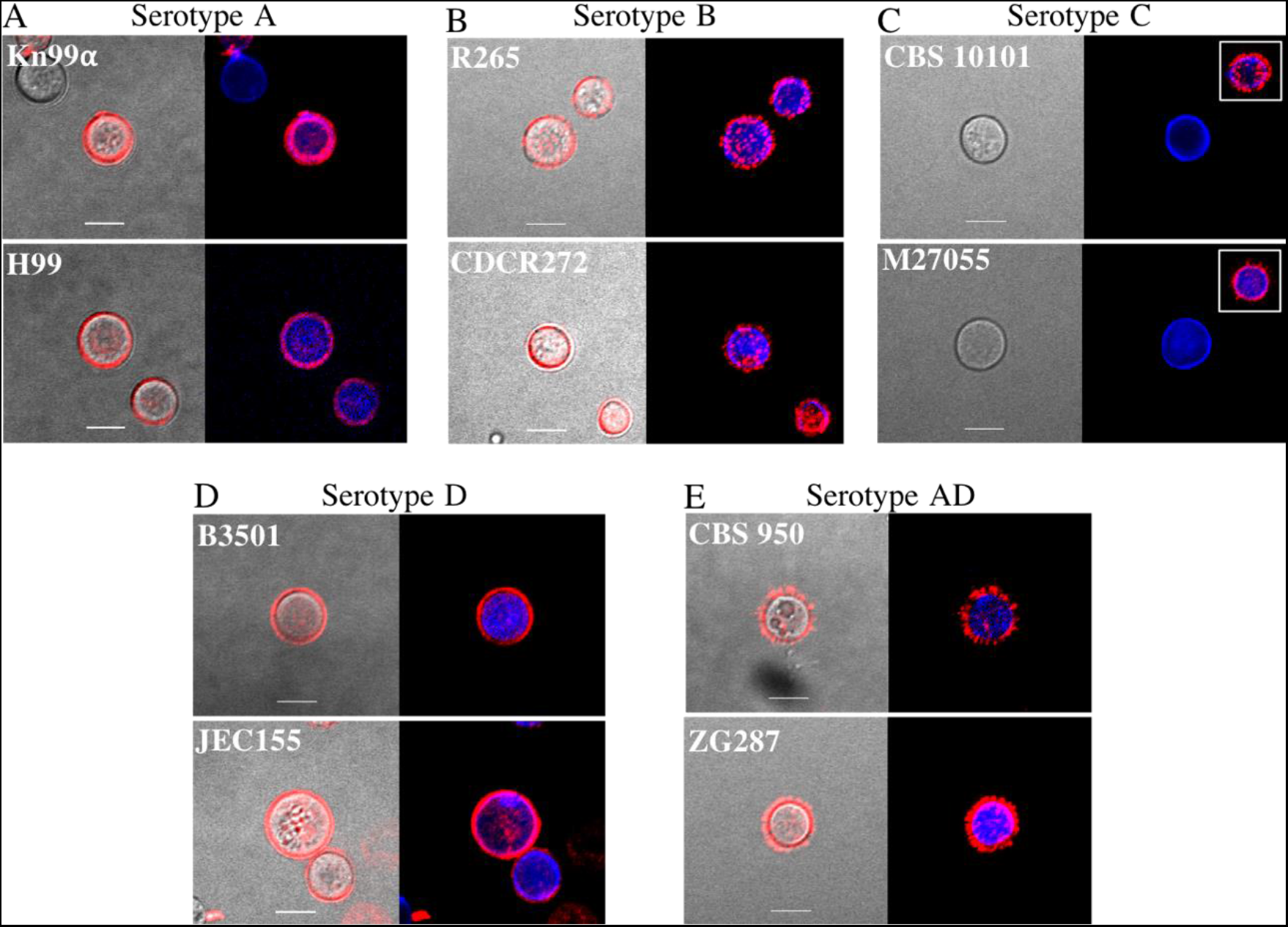
The immunofluorescence-binding pattern of Crp127 correlates with serotype. Two Cryptococcus strains from **A)** serotype A (Kn99 and H99), **B)** serotype B (R265 and CDCR272), **C)** serotype C (CBS 10101 and M27055), **D)** serotype D (B3501 and JEC155) and **E)** serotype AD (CBS 950 and ZG287) were labelled for chitin (blue; CFW) and Crp127 (far-red; goat Alexa-647-conjugated anti-mouse IgM μ-chain) and maximum-intensity projections generated via confocal microscopy. Representative images are shown for each strain. Images are merged for transmitted light and Crp127 (left panels) and Crp127 and chitin (right panels). Insets in the top right of images show representative cells labelled for chitin (blue) and O-acetyl-independent mAb F12D2 (far-red; Alexa-647-conjugated F(abߣ)2 goat anti-Mouse IgG (H+L)). Scale bars represent 5 μm.

As annular and punctate binding patterns have been associated with opsonic and non-opsonic anti-GXM IgM mAbs, respectively, we tested the ability of Crp127 to opsonise cells from strains Kn99α (annular) and R265 (punctate). Unlike positive control treatments mAb 18B7 and pooled human serum, Crp127 did not enhance phagocytosis of either strain by J774 macrophage-like cells in the presence or absence of serum (fig. S3). In summary, annular binding patterns are associated with the high-level binding of Crp127 to *C. neoformans* serotypes A and D strains. On the other hand, punctate binding is associated with low-level binding of Crp127 to serotype B strains. However, under the conditions tested in this study, neither binding pattern is clearly associated with opsonic efficacy.

### Crp127 epitope recognition reflects serotype differences within *C. gattii*

Our data above indicate that Crp127 binding accurately reflects known serotyping of cryptococcal strains. However, recent genomic data indicate that *C. gattii* may in fact be composed of several cryptic species (37). We therefore extended our analysis of this species group by investigating a further four *C. gattii* strains, representing molecular subtypes VGI-VGIII. Similar levels of Crp127 epitope recognition was seen for serotype B strains R265 (VGIIa), CDCR272 (VGIIb), EJB55 (VGIIc) and CA1873 (VGIIIa) (fig. 5A-B; P > 0.05), however significantly higher recognition was seen for the serotype B strain DSX (VGI) (fig. 5A-B; P < 0.01). Indirect immunofluorescence corroborated these findings, with punctate binding seen for the four strains presenting the epitope at low levels (fig. 5D-H) and annular binding seen for strain DSX (fig. 5C). We also tested strain CA1508 (VGIIIb), a *C. gattii* strain that, to our knowledge, has not previously been serotyped. Both flow cytometry and indirect immunofluorescence showed that Crp127 did not recognise this strain (fig. 5A and H), implying that it is a serotype C strain. In combination with the data presented in figure 3, our finding that four out of five serotype B strains were bound similarly by Crp127 suggests that availability of this epitope is fairly well conserved within this serotype.

**Fig. 5.**
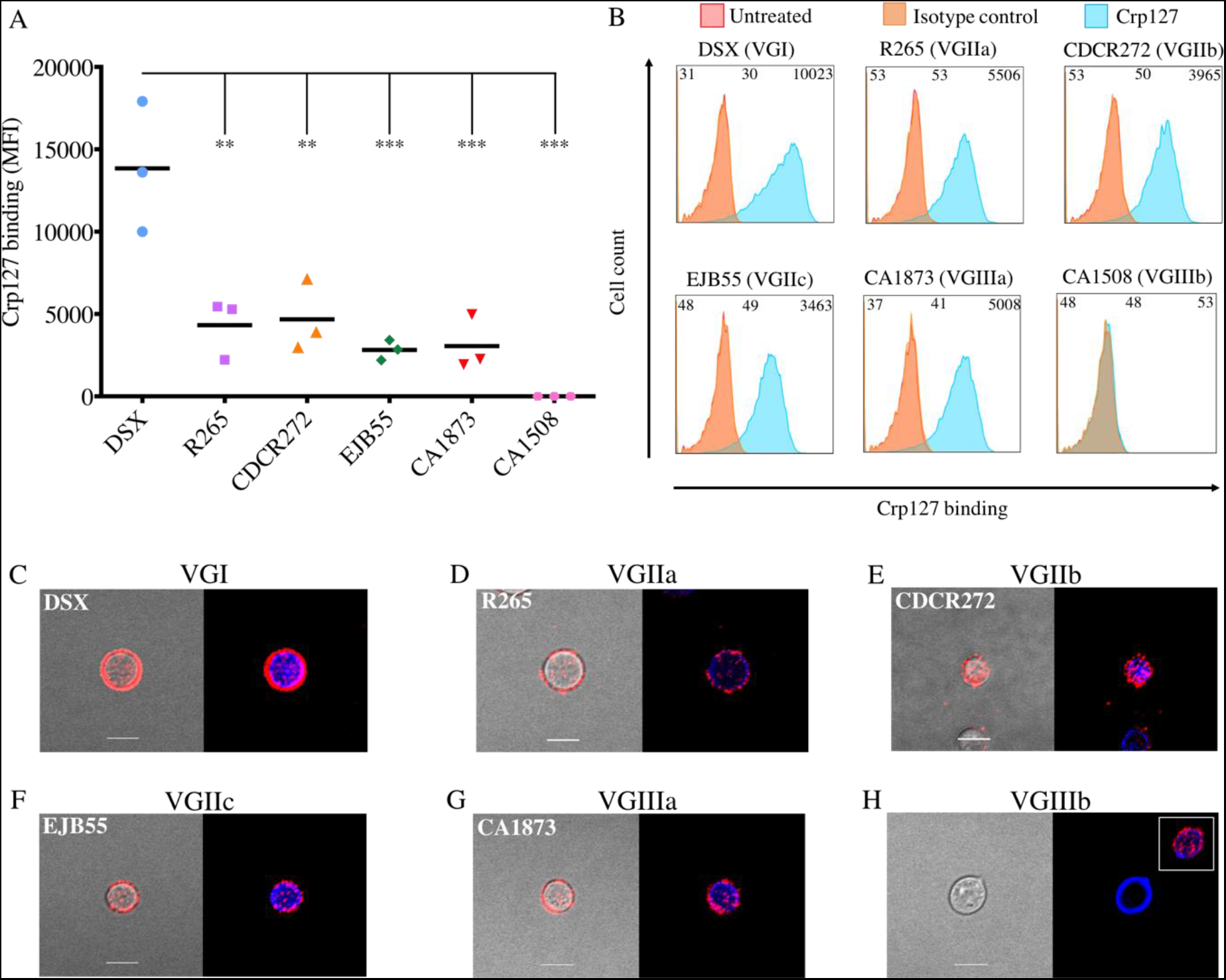
Recognition of the Crp127 epitope is largely consistent within C. gattii serotypes. The ability of Crp127 to bind to six strains from C. gattii that encompass molecular types VGI-VGIIIb was quantified via flow cytometry. **A)** Scatter plots show corrected MFI values for each strain, which were calculated by subtracting the MFI value of isotype control cells from the MFI value of the corresponding Crp127-treated cells. Data points represent MFI values calculated from three biological replicates performed as independent experiments. Tukey’s multiple comparisons test was used to test the statistical significance of differences between the six strains (n=3) (ns P > 0.05; ** P < 0.01; *** P < 0.001). **B)** Histograms show a representative distribution of Crp127 binding for strains DSX (VGI), R265 (VGIIa), CDCR272 (VGIIb), EJB55 (VGIIc), CA1873 (VGIIIa) and CA1508 (VGIIIb). Fluorescence intensity values for untreated, isotype control and Crp127-treated cells are presented in red, blue and orange, respectively, with corresponding MFI values displayed in the top left, centre and right of each panel. C. gattii strains **C)** DSX, **E)** R265, **F)** CDCR272**, G)** CA1873 and **H)** CA1508 were labelled for chitin (blue; CFW) and Crp127 (far-red; goat Alexa-647-conjugated anti-mouse IgM μ-chain) and maximum-intensity projections generated via confocal microscopy. Presented are representative images merged for transmitted light and Crp127 (left panels) and Crp127 and chitin (right panels). Scale bars represent 5 μm. Insets in the top right of images show representative cells labelled for chitin (blue) and O-acetyl-independent mAb F12D2 (far-red; Alexa-647-conjugated F(abߣ)2 goat anti-Mouse IgG (H+L)). Scale bars represent 5 μm.

### The Crp127 epitope localises to spatially confined zones of the enlarged capsule and binding elicits capsular swelling reactions

Having investigated the binding of Crp127 to cells with a small capsule, we next wished to investigate cells that had been grown in capsule-inducing conditions, given that capsule enlargement occurs shortly after infection of the host. Interestingly, in all of the strains tested we saw that the Crp127 epitope was spatially confined to distinct capsular regions (fig. 6). For serotype A and D strains, antibody binding was detected in the central zone of the capsule (fig. 6A and D). Serotype B strains differed, with regions adjacent to the cell wall and on the capsule surface bound by Crp127 in the case of strain R265 but only the single region proximal to the surface bound in the case of CDCR272 (fig. 6B). Serotype AD strain ZG287 exhibited a similar pattern of binding to R265, with Crp127 binding to both an inner and outer region of the capsule, however strain CBS 950 was bound in a region adjacent to the cell wall (fig. 6D).

The binding of mAbs to capsular GXM alters the refractive index of the enlarged capsule, resulting in capsular swelling reactions that can be visualised using DIC microscopy (38). In testing the ability of Crp127 to produce a capsular swelling reaction with strains Kn99α, R265, B3501 and CBS 950, we observed no discernible differences in the reaction pattern produced between strains, with a highly refractive outer rim and a textured inner capsule characteristic for each strain (fig. 6E-H; left panels). Notably, however, Crp127 reaction patterns differed from those elicited by 18B7, which also exhibited a highly refractive outer rim but lacked texture throughout the capsule (fig. 6E-H; right panels). Taken together, our studies of Crp127 binding to capsule-induced cells demonstrate that the Crp127 epitope is localised to specific capsular regions and that Crp127 binding produces capsular swelling reactions that are independent of serotype.

**Fig. 6.**
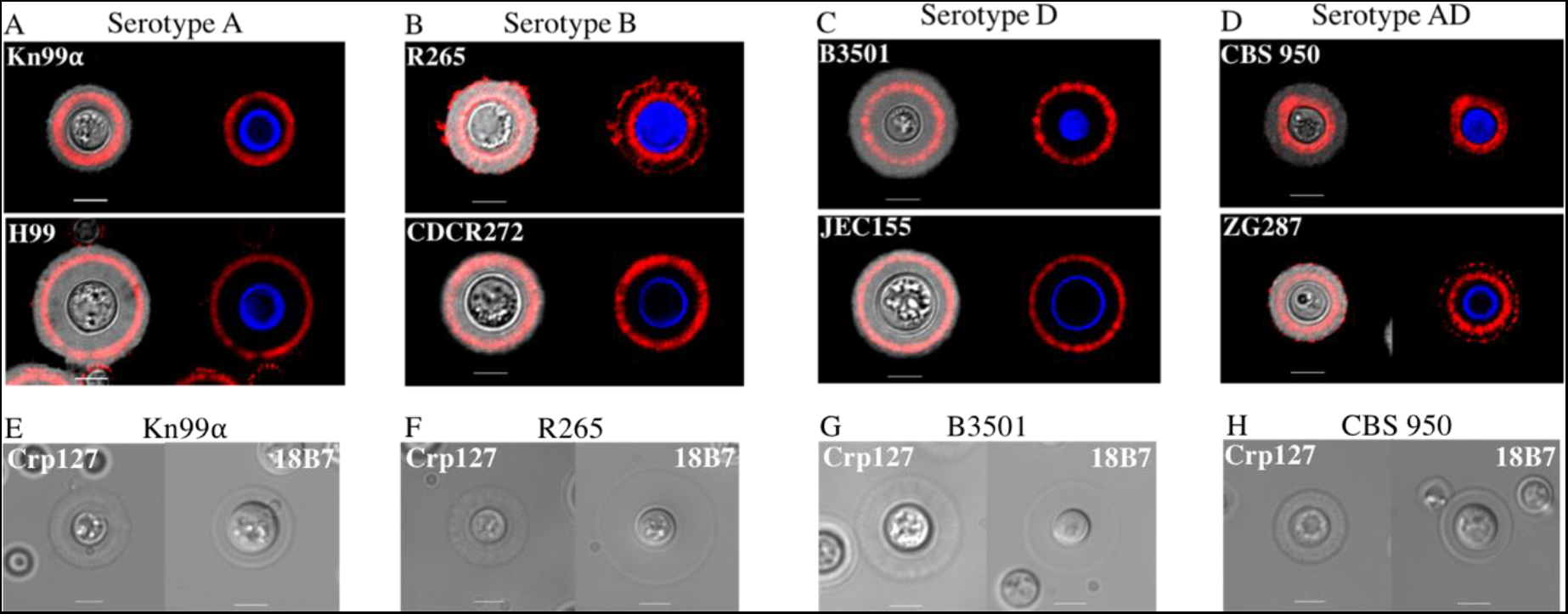
The Crp127 epitope is spatially confined to distinct capsular regions and binding elicits capsular swelling reactions distinct from those of 18B7. Cryptococcus cells were grown in capsule-inducing conditions and imaged to determine the location of the Crp127 epitope within the enlarged capsule and characterise the capsular reaction patterns elicited by this antibody. Cryptococcus strains from **A)** serotype A (Kn99α and H99), **B)** serotype B (R265 and CDCR272, **C)** serotype D (B3501 and JEC155) and **D)** serotype AD (CBS 950 and ZG287) were labelled for chitin (blue; CFW) and Crp127 (far-red; goat Alexa-647-conjugated anti-mouse IgM μ-chain), suspended in Indian ink to visualise the capsule and imaged using confocal microscopy. Representative images of a single focal plane are shown for each strain. Presented are images merged for transmitted light and Crp127 (left panels) and Crp127 and chitin (right panels). Capsule-induced cells of strains **E)** Kn99α, **F)** R265, **G)** B3501 and **H)** CBS 950 were also treated with either Crp127 (left panels) or 18B7 (right panels) and imaged using DIC microscopy to observe capsular reaction patterns. Scale bars represent 5 μm.

### Spatial distribution of the Crp127 epitope differs within the capsules produced by Titanide, yeast-like and Titan cells

Following infection of the host lung, a proportion of *C. neoformans* cells differentiate into Titan cells, a very large morphotype that facilitates pathogenesis and is associated with poor clinical outcomes (8, 12). When grown under Titanising conditions *in vitro*, *C. neoformans* forms a heterogeneous population of small oval-shaped Titanide cells (<5 μm cell body diameter), yeast-like cells (~5 μm) and large Titan cells (>10 μm) (9). As differences in capsule structure – but not mAb binding pattern – are known to exist between yeast and Titan cells (26, 39), we tested whether Crp127 could distinguish the morphological subtypes found in Titanising populations from strains H99 and Kn99α. Indeed, when imaging cells grown in Titanising conditions *in vitro*, we noticed differences in the spatial distribution of the Crp127 epitope within the capsules produced by cells of different sizes (fig. 7A). Specifically, Crp127 bound to a capsular region adjacent to the cell wall in smaller Titanide cells, within the central zone of the capsule in yeast-like cells and close to the capsule surface of Titan cells. In order to quantify how cell size affects capsular distribution of the Crp127 epitope, we determined the ratio between the area of capsule encompassed by the Crp127 epitope and the area of the whole capsule; using this metric, a ratio approaching 1 is indicative of the epitope being found in close proximity to the capsule surface (fig. 7B). Across three biological repeats (with a mean number of 111 and 133 cells measured for strains H99 and Kn99α, respectively), mean (± standard error of the mean) ratios of 0.28 ± 0.04 and 0.30 ± 0.03 were calculated for Titanide cells (<5 μm in diameter) for strains H99 and Kn99α, respectively, consistent with our initial observations that the Crp127 bound near to the cell wall of these small cells (fig. 7C-D). For yeast-like cells (5–10 μm in diameter), mean ratios were 0.42 ± 0.03 and 0.39 ± 0.02 for strains H99 and Kn99α, respectively (fig. 7C-D), indicating the Crp127 epitope is predominantly located in the central zone of the capsule in yeast-like cells, as we had previously observed (fig. 6A). Finally, the mean ratios for Titan cells (>10 μm in diameter) were 0.72 ± 0.03 and 0.71 ± 0.03 for strains H99 and Kn99α respectively, making them significantly higher than those calculated for both Titanide (fig. 7C-D; H99 P < 0.001; Kn99α P < 0.001) and yeast-like cells (fig. 7C-D; H99 P < 0.01; Kn99α P < 0.001). In summary, our results demonstrate that Crp127 binds closer to the capsule surface of Titan cells than Titanide and yeast-like cells.

**Fig. 7.**
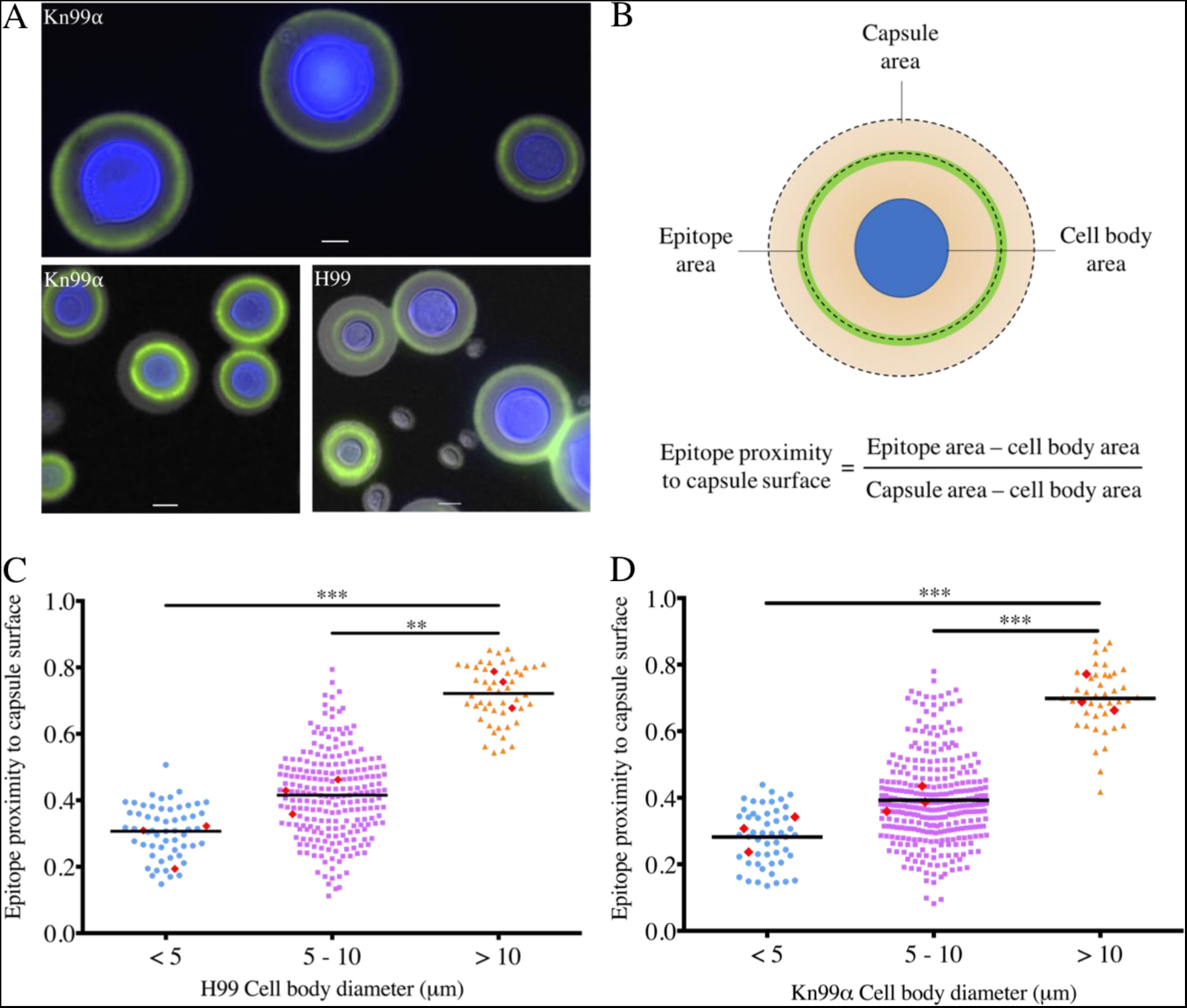
Spatial distribution of the Crp127 epitope differs within the capsule of the three cell subtypes found in Titanising populations. Cultures of C. neoformans strains H99 and Kn99α that derived solely from Titan cells were investigated for differences in the capsular distribution of the Crp127 epitope. **A)** Representative images of cells from strains H99 and Kn99α grown under conditions permissive for Titanisation. Scale bars represent 5 μm. **B)** A schematic representation of how Crp127 epitope proximity to the capsule surface was quantified through the analysis of micrographs using ImageJ software. The proximity of the Crp127 epitope to the capsule surface of **C)** H99 and **D)** Kn99α cells with cell body diameter <5 μm (blue circles), 5–10 μm (purple squares) and >10 μm (orange triangles) was quantified. Data points represent all individual cells for which the location of the Crp127 epitope was quantified, whilst the horizontal bar represents the mean. Red diamonds represent mean values calculated from three biological repeats. Tukey’s multiple comparisons test was used to test for statistically significant differences between the three groups (n=3) (** P < 0.01; *** P < 0.001).

### Migration of the Crp127 epitope towards the surface of the capsule coincides with cell enlargement

To investigate the effect of small changes in cell size on Crp127 epitope distribution, we plotted cell body diameter against epitope proximity to the capsule surface for all cells measured (fig. 8A-B). Having done so, we noticed a correlation between the cell body diameter and epitope proximity to the capsule surface of yeast-like cells. In agreement with this, when plotting only cells with a cell body diameter between 5–10 μm, we found a positive correlation between cell body diameter and epitope proximity to the capsule surface in both strains tested (fig. 7F-G; H99 r = 0.65; Kn99α r = 0.66). Unlike cell body diameter, capsule diameter did not correlate with epitope proximity to the capsule surface, indicating that changes in capsule size do not explain changes in the proximity of the Crp127 to the capsule surface (fig. S4).

**Fig. 8.**
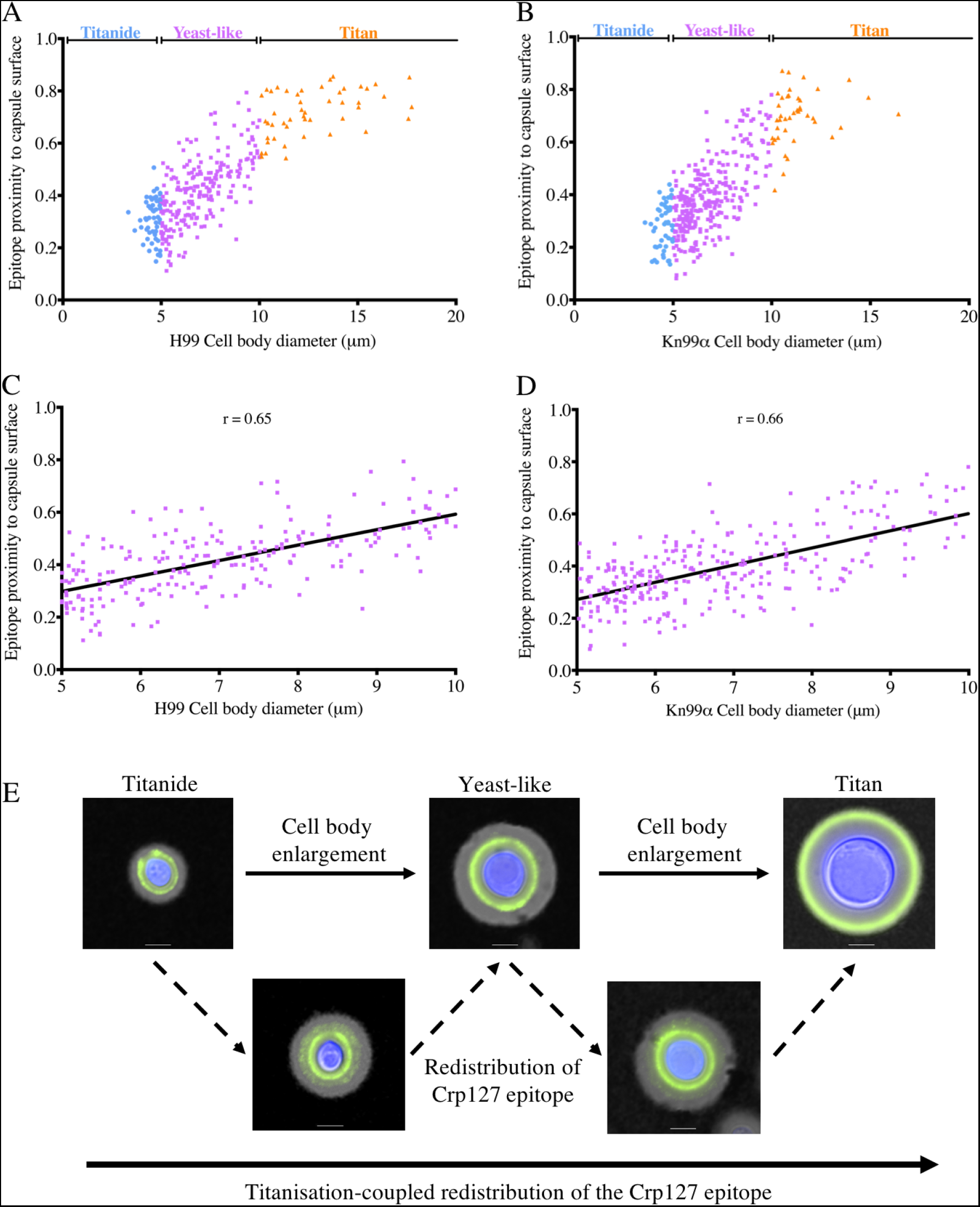
Cell size positively correlates with proximity of the Crp127 epitope to the capsule surface and suggest a model of Titanisation-coupled epitope redistribution. Cell body diameter was plotted against the epitope proximity to the capsule surface for all cells measured from strains **A)** H99 and **B)** Kn99α. Data points represent individual cells with cell body diameter <5 μm (blue circles), 5–10 μm (purple squares) and >10 μm (orange triangles). Linear regression analysis of cells with cell body diameter between 5–10 μm was used to assess the correlation between cell size and Crp127 epitope proximity to the capsule surface for strains **C)** H99 and **D)** Kn99α. **E)** Positive correlation between the cell body diameter and proximity of epitope to the capsule surface suggests a model by which Titanisation is coupled to redistribution of the Crp127 epitope towards the capsule surface. Representative images of Titanide, yeast-like and Titan cells exhibiting a typical annular immunofluorescence binding pattern – in addition to Titanide and yeast-like cells exhibiting a second more faint ring of antibody binding pattern outside of this – are presented to propose a model by which epitope is added closer to the capsule surface as cells enlarge. Images are merged for transmitted light, and Crp127 (green) and chitin (blue). Scale bars represent 5 μm.

Our results suggest that the Crp127 epitope moves gradually to the capsule surface as cells enlarge and age, raising the question of how this may occur. Throughout our imaging experiments, the binding of Crp127 to the majority of Titanide and yeast-like cells (in addition to all Titan cells) produced an annular immunofluorescence binding pattern (fig. 8E). However, we also noticed that some Titanide and yeast-like cells produced a second more faint ring of Crp127 epitope outside of this typical annular ring (fig. 8E). This may represent the addition of Crp127 epitope closer to the capsule surface, partially explaining how redistribution of this epitope coincides with cell enlargement.

## Discussion

In this study, we demonstrated that a capsular epitope recognised by Crp127 – an anti-GXM mAb produced in our laboratory – contributes to serotype-specific differences in capsule structure. This epitope traverses the capsule as cells enlarge under conditions permissive for Titanisation, resulting in its differential distribution throughout the capsule of the three *C. neoformans* morphotypes found within Titanising populations. Detailing the accessibility and localisation of this epitope adds to the existing body of literature surrounding the variability of the cryptococcal capsule between strains and reveals yet another way in which Titan cell capsules are structurally distinct from those produced by yeast cells (21–23, 32, 40).

Based on our examination of a panel of mutants harbouring capsule defects, we propose that Crp127 is an anti-GXM mAb whose binding depends on GXM *O*-acetylation, but not xylosylation. When comparing sequences of the CDRs from Crp127 with four previously characterised anti-GXM IgM mAbs – namely 2D10, 12A1, 13F1 and 21D2 – we found that Crp127 CDRs were significantly different, particularly with regard to the light-chain variable (V_L_) CDRs. These differences reflect differential gene usage and are likely to manifest as differences in epitope specificity (41, 42). In contrast, when we aligned the heavy-chain variable (V_H_) and V_L_ sequences from Crp127 with those from anti-GXM IgG1 mAb 302, we noticed that the sequences were extremely similar as a result of identical variable region gene segment usage by these two mAbs. Identical gene segment usage is not entirely surprising given the restricted set of antibody gene segments utilised by antibodies specific to capsular polysaccharides (29), however the two mAbs were produced in response to GXM derived from different serotypes of *Cryptococcus*. Whereas mAb 302 was generated following the immunisation of a mouse with serotype D GXM (ATCC 24064) (43), we generated Crp127 through the immunisation of a mouse with a cocktail containing both serotype A (H99) and serotype B (R265) GXM. Whichever serotype of GXM activated the B cell from which Crp127 derives, the sequence similarities between mAbs Crp127 and 302 demonstrate that nearly identical antibodies can be elicited during infection by at least two different serotypes of *Cryptococcus*.

Crp127 binding shows strong serotype dependence, with serotype D strains being recognised most strongly, followed by serotype A strains. *C. gattii* serotype B strains show lower, heterogeneous Crp127 epitope recognition and a punctate immunofluorescence binding pattern, whilst serotype C strains entirely fail to bind the antibody. Interestingly, the predominant SRG found in GXM produced by serotype D, A, B and C contains 1, 2, 3 and 4 xylose substituents, respectively (15, 44). Together with the previous observation that β-(1,2)-xylose and *O*-acetyl groups are not added to the same backbone mannose residue (17, 36), this differential SRG usage may explain the variable Crp127 epitope recognition in one of two ways. For example, the additional xylose substituents present in the predominant SRG found in serotype B and C GXM may prevent addition of *O*-acetyl groups in such a way that the Crp127 epitope is not formed. Alternatively, the extra xylose substituents found in these SRGs may sterically hinder binding of Crp127 to its epitope. Studies that further elucidate the roles of specific proteins in GXM biosynthesis – together with advances in techniques that enable chemical synthesis of GXM oligosaccharides – will enhance our understanding of molecular how epitope recognition is achieved by anti-GXM mAbs like Crp127. Intriguingly, a recent transcriptomics study identified *CAS31* as being absent from the genome of strain CBS 10101, a serotype C isolate that we subsequently found was not recognised by Crp127. Whilst we cannot rule out the possibility that other factors contribute to the inability of Crp127 to recognise serotype C strains, it is tempting to speculate that the loss of *CAS31* function in this lineage may explain its lack of reactivity with Crp127 (32, 34).

Perhaps our most striking observation regarding the Crp127 epitope was its differential distribution throughout the capsules produced by Titanide, yeast-like and Titan cells. No such differences have been consistently observed for other anti-GXM mAbs E1, 2D10, 13F1 or 18B7 (7, 26), making this a unique feature of Crp127. The positive correlation between cell size and Crp127 epitope proximity to the capsule surface is suggestive of a scenario whereby the epitope in question is initially produced in a capsular region adjacent to the cell wall in small Titanide cells, before redistributing first to the midzone of yeast-like cells and eventually to the capsule surface of Titan cells. This finding raises the intriguing question of how formation and removal of the Crp127 epitope is so tightly spatially controlled within the capsule? One possibility is that the epitope could be formed at the cell surface and then move outwards as the capsular material elongates. Therefore, we speculate that since the epitope moves outwards at a faster rate than the capsule expands, and since the amount of epitope that initially surrounds a smaller Titanide or yeast-like cell would not be sufficient to form the perimeter of capsule encasing a much larger Titan cell, we instead favour a model in which the epitope is enzymatically removed and added to different regions of the capsule during growth. For instance, it is possible that GXM decorated with *O*-acetyl groups is added closer to the capsule surface in larger cells or that such regions are “unmasked” in a different capsular region as the capsule is reshaped during Titanisation (26). To summarise, our findings demonstrate that the differential distribution of specific epitopes within the cryptococcal capsule is yet another way in which Titan cells can be distinguished from canonical yeast cells, prompting further investigation into how redistribution of such epitopes occurs and what impact this has on the outcome of infection.

## Materials and methods

### Reagents, strains and mAbs

All reagents were purchased from Sigma-Aldrich unless stated otherwise. The *Cryptococcus* strains used in this study are described in table S1. The anti-GXM mAbs used in this study are described in table S2.

### Growth of cryptococci

*Cryptococcus* strains were preserved at −80°C in MicroBank_TM_ tubes (Thermo Fisher Scientific) prior to being stored on yeast extract peptone dextrose (YPD) agar plates at 4°C for a maximum of 30 days. Unless stated otherwise, strains were cultured on a rotary wheel at 20 rpm for 24 h at 25°C in round-bottom culture tubes containing 3 mL YPD broth. To induce capsule growth, *Cryptococcus* cells were grown in round-bottom culture tubes containing 3 mL Dulbecco’s Modified Eagle’s Media (DMEM) supplemented with 2 mM L-glutamine, 100 U/mL penicillin, 100 U/mL streptomycin and 10% foetal bovine serum (FBS) for 72 h in an incubator at 37°C and 200 rpm.

### Hybridoma production and mAb purification

Cultures of *C. neoformans* H99 and *C. gattii* R265 were microfuged (4000 × *g* for 5 min) and washed three times in 1 mL Dulbecco’s phosphate-buffered saline (PBS). Washed cultures were then heat killed for 60 minutes at 65°C. Following heat killing 20 μL was plated onto YPD to confirm there were no viable cells. Heat-killed H99 and R265 cells were then either lysed (see below) or mixed 1:1 and stored at −20°C prior to inoculation. Fungal cells were lysed using Precellys tubes (UK05 03961-1-004) using programme 6400-2×10-005. Following lysis, lysis beads were microfuged (3000 × *g* for 1 min) and supernatant collected. H99 and R265 lysates were mixed 1:1 and stored at −20°C.

BALB/c mice were hyper-immunised with heat-killed H99 and R265 in addition to their lysates. Hybridomas were generated by a method that has previously been described (45). NS0 immortal fusion partner cells were fused with splenocytes mediated by polyethylene glycol (StemCell Technologies). All animal work was conducted in accordance with Home Office guidelines and following local ethical approval granted under animal licence 30/2788. Supernatants from clones were screened for reactivity with H99 and R265 cells using 96-well plates, with FITC-conjugated anti-mouse IgG and anti-mouse IgM antibodies used to identify positive clones via epifluorescence microscopy. Positive clone 127 was cultured in RPMI 1640 with IgG-depleted FBS and supernatant collected in a MiniPerm bioreactor (Sarstedt). MAb Crp127 was purified from supernatant using affinity chromatography and ProSep Thiosorb (Millipore).

### Hybridoma sequencing and antibody sequence analysis

Sequencing of hybridomas was carried out by Absolute Antibody Ltd (UK). Sequencing was performed by whole transcriptome shotgun sequencing (RNA-Seq). In brief, hybridomas were cultured in Iscove’s Modified Dulbecco’s Media (IMDM) supplemented with 10% FBS in an incubator at 37°C and with 5% CO_2_. Total RNA was extracted from cells and a barcoded cDNA library generated through RT-PCR using a random hexamer. Sequencing was performed using an Illumina HiSeq sequencer. Contigs were assembled and annotated for viable antibody sequences (i.e those not containing stop codons) to confirm the species and isotype of mAb Crp127 as murine and IgM, respectively.

Variable region gene usage was determined using VBASE2 software (46) and CDRs were predicted using the Kabat numbering system (47). Heavy-chain variable (V_H_) and light-chain variable (V_L_) sequences of mAb Crp127 were aligned with antibody sequences that have previously been described (35, 48). Amino acid sequences were aligned using Clustal Omega software (49) and annotated using ESpript software (50).

### Immunolabelling

*Cryptococcus* cells were immunostained for flow cytometry and microscopy experiments. 1 mL of fungal culture was transferred to a 1.5 mL microcentrifuge tube, microfuged (15,000 × *g* for 1 min) and washed 3x in PBS. Cell density was determined using a haemocytometer and adjusted to 10_7_ cells/mL in a final volume of 200 μL. 20 μg/mL Crp127, F12D2, 18B7 or mouse anti-human IgG (IgM isotype control) were added and samples mixed on a rotary wheel at 20 rpm for 1 h at room temperature. Untreated cells for use in flow cytometry were left untreated. After primary antibody treatment, samples were microfuged (15,000 × *g* for 1 min) and washed 3x in PBS to remove unbound primary antibody. 2 μg/mL Alexa-488-conjugated goat anti-mouse IgM (heavy chain) (Thermo Fisher Scientific), Alexa-647-conjugated goat anti-mouse IgM μ-chain (Abcam) or Alexa-647-conjugated F(abߣ)2-Goat anti-Mouse IgG (H+L) (Thermo Fisher Scientific) were added to antibody-treated samples and samples mixed on a rotary wheel at 20 rpm for 1 h at room temperature. Secondary antibody was also added to isotype control samples for flow cytometry. For microscopy experiments, 5 μg/mL calcofluor-white (CFW) was also added at this stage to label chitin. Following incubation with secondary antibody, samples were again microfuged (15,000 × *g* for 1 min) and washed 3x to remove unbound secondary antibody and CFW.

### Flow cytometry

Flow cytometry experiments were performed with an Attune NxT Flow Cytometer equipped with an Attune Autosampler (Thermo Fisher Scientific). Untreated, isotype control and either Crp127 or 18B7 samples were prepared for each strain or conditions tested. Following immunostaining, samples were diluted to 5 × 10_6_ cells/mL and 200 μL of *Cryptococcus* put into individual wells of a plastic round-bottom 96-well plate ready for insertion into the Attune Autosampler. Sample was collected from each well at a rate of 100 μL/min until 10,000 events were recorded. The 488 nm laser was used to detect primary antibody bound by Alexa-488-conjugated secondary antibodies, with the same voltage used to power the laser within each experiment. Flow cytometry data was then analysed using FlowJo (v10) software. Debris was excluded by using the FSC-A vs. SSC-A gating strategy, followed by exclusion of doublets using the FSC-A vs. FSC-H gating strategy (fig. S5). Where GXM-deficient mutants were analysed, samples were only gated to exclude debris due to the inseparable large aggregates formed by these mutants as a result of budding defects. After gating, histograms of fluorescence intensity were plotted and the median fluorescence intensity (MFI) determined. Corrected MFI values were calculated by subtracting the MFI value of the mAb-treated sample by the corresponding isotype control sample in the case of Crp127 or untreated sample where 18B7 was used. Across all experiments, MFI values returned from isotype control cells were extremely similar to those returned from untreated cells.

### Confocal microscopy

Following the final washes of the immunostaining procedure, 2 μL of stained cryptococcal cells were spotted onto a glass slide and placed under a square glass coverslip. Where visualisation of the capsule was necessary, 2 μL Indian ink was also added to the glass slide. Imaging was performed on a Nikon A1R laser scanning confocal microscope using a 100x object lens and oil immersion. Alongside transmitted light, 639 nm and 405 nm lasers were used to detect Alexa-647-conjugated secondary antibodies and CFW, respectively. For cells with small capsules, Z-stacks spanning 8 μm were generated using steps of 0.27 μm. For capsule-induced cells, Z-stacks were taken across 20 μm using steps of 0.66 μm. Generation of maximum intensity projections (MIPs) and other image processing was performed using NIS-Elements and ImageJ software.

### Chemical de-*O*-acetylation of capsular GXM

Where chemical de-*O*-acetylation of the capsule was required, cells were grown in YPD broth that had been adjusted pH 11 with NaOH and sterilised with a 0.22 μm filter. Round-bottom culture tubes containing 3 mL of pH 11 YPD broth was then placed on a rotary wheel turning at 20 rpm for 24 h at 25°C. This method was adapted from that used in a previous study (21).

### Phagocytosis assays

Phagocytosis assays were performed using the murine macrophage-like J774A.1 cell line (mouse BALB/cN; ATCC_®_ TIB-67™). Cells were cultured in DMEM supplemented with 2 mM L-glutamine, 100 U/mL penicillin, 100 U/mL streptomycin and 10% FBS, before 1 × 10_5_ cells were seeded onto round glass coverslips that had been placed into wells of a flat-bottom 24-well plate and incubated for 24 h at 37°C and 5% CO_2_. Cells of strain R265 and Kn99α were opsonised with 18B7 or Crp127 as described for the first incubation of the immunostaining procedure. In the same way, cells were opsonised with 10% AB-human serum alone or in combination with Crp127. To achieve a multiplicity of infection (MOI) of 10, 10_6_ R265 or Kn99α cells were then resuspended in serum-free DMEM and added to each well of J774A.1 cells. Following the infection, each well was gently washed 3x with 1 mL of warmed PBS to remove extracellular yeast. The contents of each well were then fixed with 4% paraformaldehyde prior to being washed a further 3x. Cover slips were then extracted from their well, any residual PBS removed by briefly submersing in sterile dH_2_O and mounted onto glass slides with Prolong Gold Antifade Mountant (Thermo Fisher Scientific). The total number of internalised yeast cells per 100 J774A.1 cells (phagocytic index) was determined by microscopic examination using a Nikon TE2000-U microscope with a 60x objective lens and oil immersion.

### Capsular swelling reactions

Capsule-induced cells were treated with 50 μg/mL Crp127 or 18B7 as described for the immunostaining procedure. 2 μL of *Cryptococcus* cells were then dropped onto a glass slide and placed under a square glass coverslip. Imaging was performed on the differential interference contrast (DIC) channel of a Nikon TE2000-U microscope using a 60x objective lens with oil immersion. Image processing was performed using NIS-Elements and ImageJ software.

### Titan cell experiments

Titan cells that exhibit all the properties of *in vivo* Titan cells were induced *in vitro* using a previously described protocol (9). *C. neoformans* H99 and Kn99α cells were cultured in glass conical flasks containing 10 mL yeast nitrogen base (YNB) + 2% glucose at 30°C and 200 rpm for 24 h. Cells were adjusted to an OD_600_ reading of 0.001 before being transferred into 10% heat-inactivated foetal calf serum (HI-FCS) at a final volume of 3 mL in a plastic six-well plate and grown for 72 h at 37°C and 5% CO_2_. To begin a culture derived solely from Titan cells, cells were passed through an 11 μm filter, trapping only larger cells on the filter paper. This filter paper was then washed in PBS to re-suspend Titan cells. Between 10_3_ and 10_4_ Titan cells were then transferred into 3 mL HI-FCS in a plastic six-well plate and cultured for a further 72 h at 37°C and 5% CO_2_. Titanising populations were prepared for imaging according to the method described for immunostaining. Imaging was performed on a Nikon TE2000-U microscope using a 60x objective lens with oil immersion.

To quantify the proximity of the Crp127 epitope to the capsule surface was quantified, ImageJ software was used to draw regions of interest (ROIs) around the cell body, the immunofluorescence binding pattern of the Crp127 and the capsule surface (as determined by Indian ink staining). For each cell measured, the area of these three ROIs was determined before the area of the cell body was subtracted from the areas calculated for both the Crp127 epitope ROI and the capsule surface ROI. Finally, the area of the Crp127 epitope ROI was divided by the capsule surface ROI as a means of quantifying the proximity of the Crp127 epitope to the capsule surface. A mean number of 111 and 133 cells were measured per biological replicate for strain H99 and Kn99α, respectively. Image processing was performed using NIS-Elements software.

### Experimental design and statistical analysis

For each experiment described, three biological repeats were performed as independent experiments that were carried out on different days. All datasets were analysed using GraphPad Prism 7 software.

## Acknowledgements

**Provision of antibodies, strains and assistance**

We gratefully acknowledge our colleagues Tamara Doering (Washington University), Guilhem Janbon (Institut Pasteur), Arturo Casadevall (Johns Hopkins) and Thomas Kozel (University of Nevada) providing antibodies and strains and for their invaluable advice regarding this project. We are also grateful to Alessandro Di Maio, Leanne Taylor-Smith and Joao Correia (University of Birmingham) for assistance with confocal microscopy and subsequent image processing.

## Author contributions

Experiments were designed and conducted by MP, XZ and EB. The Crp127 antibody was raised and initially characterised by SAJ and MG. ERB and RCM helped design and oversee this project. Data figures and text were prepared by MP and then edited and revised by all the other authors.

## Funding

This work was made possible via funding from the Lister Institute for Preventive Medicine and the European Research Council under the European Union’s Seventh Framework Programme (FP/2007-2013)/ERC Grant Agreement No. 614562 and from the Biotechnology and Biological Sciences Research Council (BBSRC) via grant BB/R008485/1. RCM is additionally supported by a Wolfson Royal Society Research Merit Award. ERB was supported by the UK Biotechnology and Biological Research Council (http://www.bbsrc.ac.ukhttp://www.bbsrc.ac.uk; BB/M014525/1).

### Data and materials availability

All data needed to evaluate the conclusions drawn in this paper are present in the paper and/or the supplementary materials. Additional data related to this paper may be requested from the authors. The Crp127 antibody described here is available via Ximbio.com.

